# A novel strain of lumpy skin disease virus causes clinical disease in cattle in Hong Kong

**DOI:** 10.1101/2021.04.20.440323

**Authors:** John Flannery, Barbara Shih, Ismar R. Haga, Martin Ashby, Amanda Corla, Simon King, Graham Freimanis, Noemi Polo, Anne Ching-Nga Tse, Christopher J. Brackman, Jason Chan, Patrick Pun, Andrew D. Ferguson, Andy Law, Samantha Lycett, Carrie Batten, Philippa M. Beard

**Affiliations:** The Pirbright Institute, Ash Rd, Surrey, UK; The Roslin Institute, University of Edinburgh, Easter Bush, Midlothian, UK; Agriculture, Fisheries and Conservation Department, Government of the Hong Kong Special Administrative Region, Hong Kong SAR, China; CityU Veterinary Diagnostic Laboratory, City University of Hong Kong, Hong Kong SAR, China

**Keywords:** Lumpy skin disease, lumpy skin disease virus, poxvirus, cattle, Hong Kong, epidemic, phylogenetics

## Abstract

Lumpy skin disease virus (LSDV) is an emerging poxviral pathogen of cattle that is currently spreading throughout Asia. The disease situation is of high importance for farmers and policy makers in Asia. In October 2020, feral cattle in Hong Kong developed multifocal cutaneous nodules consistent with lumpy skin disease (LSD). Gross and histological pathology further supported the diagnosis and samples were sent to the OIE Reference Laboratory at The Pirbright Institute for confirmatory testing. LSDV was detected using quantitative polymerase chain reaction (qPCR) and additional molecular analyses. This is the first report of LSD in Hong Kong. Whole genome sequencing (WGS) of the strain LSDV/HongKong/2020 and phylogenetic analysis were carried out in order to identify connections to previous outbreaks of LSD, and better understand the drivers of LSDV emergence. Analysis of the 90 core poxvirus genes revealed LSDV/HongKong/2020 was a novel strain most closely related to the live-attenuated Neethling vaccine strains of LSDV and more distantly related to wildtype LSDV isolates from Africa, the Middle East and Europe. Analysis of the more variable regions located towards the termini of the poxvirus genome revealed genes in LSDV/HongKong/2020 with different patterns of grouping when compared to previously published wildtype and vaccine strains of LSDV. This work reveals that the LSD outbreak in Hong Kong in 2020 was caused by a different strain of LSDV than the LSD epidemic in the Middle East and Europe in 2015-2018. The use of WGS is highly recommended when investigating LSDV disease outbreaks.

## Introduction

Lumpy skin disease (LSD) is a severe disease of cattle and water buffalo characterised by multifocal cutaneous nodules. It is caused by infection with the poxvirus lumpy skin disease virus (LSDV), a member of the capripoxvirus genus. LSDV is transmitted by haematophagous vectors such as mosquitoes and flies which facilitates rapid spread of the virus in optimal climatic conditions [1]. LSDV is a rapidly emerging pathogen, having spread over the past ten years from Africa and the Middle East into south east Europe, the Caucasus, Russia and, more recently, Asia [2–7]. This Eurasian LSD epidemic has affected thousands of cattle and caused substantial economic loss through the loss of animals, reduced productivity, the cost of control and prevention campaigns, and loss of export markets [8]. Widespread vaccination programmes with the live-attenuated LSDV vaccine based on the Neethling strain of the virus have been key to the control of the disease in south east Europe [9, 10].

Phylogenetic analysis of LSDV isolates using whole genome sequencing (WGS) has revealed that strains sequenced to date segregate into two subgroups [11]. Subgroup 1.1 contains vaccine strains related to the original Neethling strain. Subgroup 1.2 can be divided into wildtype LSDV strains and LSDV KSGP strains. Examination of wildtype LSDV strains shows that the viral strains which caused disease in south east Europe in 2015-2016 and Russia in 2015 are very similar to a strain isolated from Israel in 2012, supporting the theory of northward spread of the virus from Africa and the Middle East. In contrast WGS of a novel strain of LSDV, known as Saratov/2017, from diseased cattle in southern Russia in 2017 revealed similarities with the Neethling vaccine strain [12]. The origin of Saratov/2017 is unclear.

We report here the clinical and pathological features of the first reported outbreak of LSD in Hong Kong in 2020, and phylogenetic analysis of the whole genome of the LSDV strain isolated from affected cattle (LSDV/HongKong/2020). This strain differs from all previously reported LSDV strains but clusters phylogenetically with Neethling vaccine strains.

## Materials and methods

Tissue samples for histopathological analysis were fixed in formalin, processed to paraffin wax blocks, sectioned, and stained with hematoxylin and eosin. EDTA blood samples (n=2), tissues (n=9), nasal swab (n=1) and serum (n=2) from three cattle were submitted to the OIE reference laboratory for LSD at The Pirbright Institute, UK, for confirmatory diagnosis. Nucleic acid was extracted from EDTA blood samples, tissues and swabs using the MagMAX™ CORE Nucleic Acid Purification Kit (ThermoFisher Scientific, Paisley, UK) on the KingFisher Flex extraction platform (ThermoFisher Scientific). Capripoxvirus DNA was detected by a qPCR assay targeting LSDV074 (also known as the p32 gene, a homolog of the vaccinia virus H3L gene) [13] using the Path-ID™ qPCR Master Mix (Thermofisher Scientific). Nucleic acid from capripoxvirus positive samples was further analysed by a qPCR targeting LSDV011 designed to differentiate the three capripoxvirus species [14] and a qPCR targeting LSDV008 designed to differentiate LSDV wildtype strains from Neethling vaccine strains [15]. Partial genome sequencing of LSDV036 (the RNA polymerase subunit RPO30) was performed using primers described by [16] and the resulting sequences were assembled using the SeqMan Pro software and phylogenetic analysis performed using MEGA7. Serum samples were tested with the ID Screen^®^ Capripox Double-antigen ELISA (ID Vet Innovative diagnostics) following the manufacturer’s instructions.

LSDV was isolated on MDBK cells (ATCC code CCL-22). Briefly, tissue homogenates from fresh skin were prepared from each of the three cattle, homogenates were sonicated twice and centrifuged prior to inoculation onto MDBK cell monolayers. Cells were harvested after 6 days and then sonicated prior to a second passage on MDBK cells using both cellular and supernatant fractions. The cells were harvested when 100% CPE was observed.

LSDV was purified as described previously [17]. Briefly, MDBK cells infected with LSDV were harvested, centrifuged at low speed, supernatants discarded and the pellets resuspended in 1 mM of Tris-HCl pH9. Samples were vortexed, sonicated and incubated with Benzonase^®^ (>250 units/ μl, Sigma E1014-25KU) ahead of two rounds of sucrose cushion purification. Samples were then treated with 33 μl of 1.5M Tris pH 8.8, 50 μl of 10% SDS, 100 μl of 60% sucrose and 85 μl of proteinase K (20 mg/ml, ThermoFisher Scientific) for 4 h at 37°C, followed by phenol-chloroform extraction and ethanol precipitation.

Total dsDNA (1 ng) was processed for WGS using the Nextera XT DNA library kit (Illumina) and performed on the Hamilton NGSStar (Hamilton Robotics). Libraries were bead normalised and pooled according to manufacturer’s protocols. Final sequencing pools were loaded at 12pM concentration onto an Illumina MiSeq 300 cycle sequencing run with a 1% PhiX spike-in (Illumina).

Using BBtools [18], adaptor and quality trimming were carried out on the reads before genome assembly. Two approaches were taken; the consensus sequence and the *de novo* assembly methods. For generating the consensus sequence, reads were mapped to the lumpy skin disease virus isolate 155920/2012 (KX894508.1) with BWA-MEM (version 0.7.17) [19], and SAMtools (version 1.10) [20] was used for processing the alignment files. The variants were detected using freebayes (version 1.3.1) [21], and the per-region coverage was estimated using mosdepth (version 0.2.6). BCFtools (version 1.10) was used for creating the consensus sequence using high quality calls (%QUAL>=20), taking into account the regions with zero coverage from the mosdepth output. *De novo* assembly was carried out using Spades (version 3.13.2) [22], setting the kmers to “33, 55, 77, 99”. Using RaGOO [23], reference-guided scaffolding was carried out on the contigs generated by SPAdes with the consensus sequence acting as the template. The consensus and *de novo* genome sequences were compared to each other using Minimap2 (version 2.16) [24]. Custom Python scripts were used to combine the two genome sequences into the final LSDV/HongKong/2020 genome sequence; in brief, where there are differences in variants, the variants in the *de novo* assembly were preferentially included, and stretches of poly-N were retained only if it was present in both the consensus and *de novo* sequences.

Prokka [25] was used for annotating the LSDV/HongKong/2020 genome, along with 16 other lumpy skin disease genomes on NCBI (MH646674.1, AF409138.1, KX764645.1, KX764644.1, MG972412.1, KX764643.1, MT134042.1, MN072619.1, AF325528.1, KX683219.1, AF409137.1, MH893760.2, KY702007.1, MN642592.1, KX894508.1, KY829023.3, see Table 1 for details). An all-against-all comparison was made on the Prokka-identified gene sequences using BLAST+ (version 2.7.1) [26]. Network analysis based on the BLAST output was performed using Graphia [27] for labelling the Prokka-identified genes with corresponding genes in the reference genome annotation (KX894508.1) in order to find a set of core genes between the LSDV genomes and to find groups of accessory genes. Each sequence (i.e. a gene in a genome) was treated as a node, two sequences were joined by an edge if they had significant similarity score (≥ 60 bitscore, ≥ 60 qcovs and evalue < 1e-5). Markov Clustering (MCL) was used for grouping nodes sharing high similarity into clusters, with each cluster containing sequences of the same gene from different genomes. In order to highlight genes that showed a strong difference between vaccine and wildtype strains, we explored the accessory gene clusters that separated to distinct components when applying an additional edge filter using 10 kNN (k-nearest neighbour) based on bitscore.

**Table 1.** Sequences included in the analysis

| GenBank ID | Abbreviated name | Year | Origin | Wildtype or vaccine | Identity to HongKong/2020 (MW732649) | Identity to vaccine (AF409138) | Identity to wildtype (KX894508) |
| --- | --- | --- | --- | --- | --- | --- | --- |
| **AF325528.1** | NI-2490 | 1959 | Kenya | wildtype | 99.54 | 99.13 | 99.93 |
| **AF409137.1** | Neethling Warmbaths LW | 1999 | South Africa | wildtype | 99.50 | 99.13 | 99.99 |
| **KX894508.1** | Israel/2012 | 2012 | Israel | wildtype | 99.50 | 99.13 | 100.00 |
| **KY702007.1** | Bunjanovac/2016 | 2016 | Bunjanovac, Serbia | wildtype | 99.50 | 99.13 | 100.00 |
| **KY829023.3** | Evros/2015 | 2015 | Evros, Greece | wildtype | 99.50 | 99.13 | 100.00 |
| **MH646674.1** | Saratov/2017 | 2017 | Saratov, Russia | wildtype | 99.59 | 99.71 | 99.40 |
| **MH893760.2** | Dagestan/2015 | 2015 | Dagestan, Russia | wildtype | 99.50 | 99.13 | 100.00 |
| **MN072619.1** | Kenya/1958 | 1958 | Kenya | wildtype | 99.53 | 99.12 | 99.92 |
| **MN642592.1** | Kubash/2016 | 2016 | Kubash, Kazakhstan | wildtype | 99.50 | 99.13 | 100.00 |
| **MT134042.1** | Udmurtiya/2019 | 2019 | Udmurtiya, Russia | wildtype | 99.57 | 99.43 | 99.63 |
| **MW732649.1** | HongKong/2020 | 2020 | Hong Kong, China | wildtype | 100.00 | 99.56 | 99.50 |
| **AF409138.1** | Neethling LW |  |  | vaccine | 99.56 | 100.00 | 99.13 |
| **KX683219.1** | KSGP 0240 |  |  | vaccine | 99.53 | 99.12 | 99.93 |
| **KX764643.1** | Neethling SIS |  |  | vaccine | 99.56 | 100.00 | 99.13 |
| **KX764644.1** | Neethling Herbivac |  |  | vaccine | 99.56 | 100.00 | 99.13 |
| **KX764645.1** | Neethling OBP |  |  | vaccine | 99.56 | 100.00 | 99.14 |
| **MG972412.1** | Neethling Croatia |  |  | Vaccine | 99.56 | 100.00 | 99.13 |

For each of the 17 genomes, the core gene sequences were merged into one continuous sequence for building a phylogenetic tree (for consecutive core genes, the intergenic regions between the genes were also included). MUSCLE (MUltiple Sequence Comparison by Log-Expectation; version 3.8.1551) [28] was used for aligning the core-gene sequences for the 17 genomes. RAxML (Randomized Axelerated Maximum Likelihood; version 8.2.12) was used to build the phylogenetic tree using a general time reversible substitution model with gamma distributed site to site rate variation (GTRGAMMA) and 100 bootstraps, which was then visualised using iTOL [29].

## Results and Discussion

There are no commercial cattle farms in Hong Kong, however there is a population of around 1000 feral brown cattle found mostly in local country parks. There is also a population of some 200 feral water buffaloes. In October 2020, multiple brown cattle in the eastern part of the New Territories of Hong Kong were reported to have multifocal nodular skin lesions (Figure 1A and B). In early November 2020 similar cases were detected in the northern New Territories close to the border with mainland China and on one of the outlying islands in the southwest. In addition to the skin lesions, other clinical signs included fever, malaise, anorexia and superficial lymphadenopathy. Nasal and/or oral ulcers were present in some cattle (Figure 1C), accompanied by nasal discharges and/or ptyalism. The clinical course lasted for 2-3 weeks and the disease was self-limiting in a majority of cases. The morbidity of the initial outbreak based on clinical signs was estimated to be between 20-30%. Two affected cattle were reported dead, however, both were over 15 years old with only mild skin lesions. It was uncertain whether death was causally associated with LSD or other unknown comorbidities. Clinical cases were absent from the buffalo populations.

**Figure 1.**
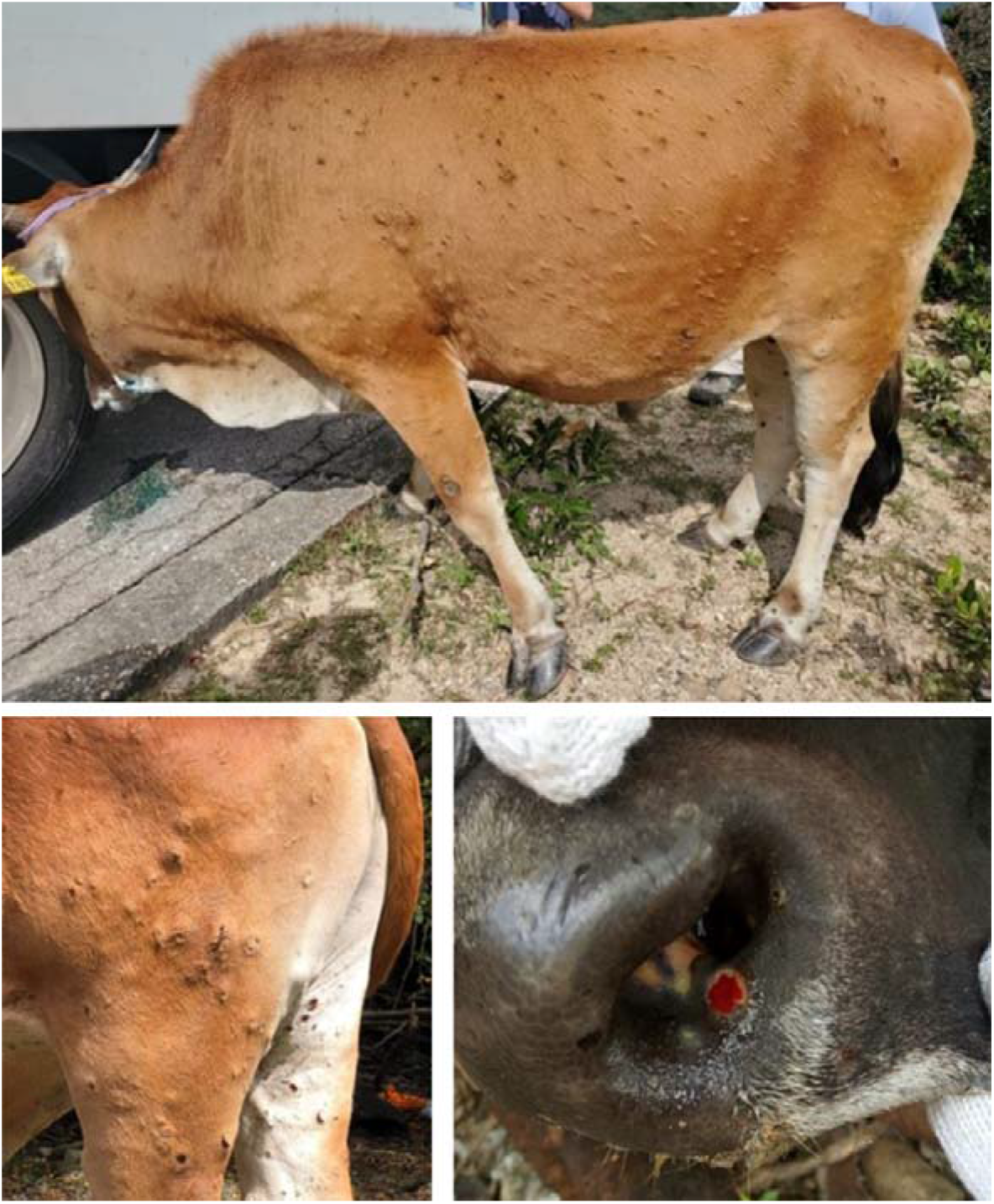
Clinical presentation of LSD in a feral brown bull, Hong Kong, 2020. (A, B) Multifocal, raised skin nodules. (C) Focal ulceration of the nasal mucosa.

Postmortem examination was carried out on two cattle at the Government’s official veterinary laboratory, Tai Lung Veterinary Laboratory. Gross and microscopic findings were consistent with a diagnosis of LSD [30]. Gross findings included widespread, randomly distributed cutaneous and subcutaneous nodules ranging from 1-40cm in diameter, sometimes with a targetoid appearance. A few nodules were ulcerated and others had a central area of dense crust (sit-fast). Multiple superficial lymph nodes were enlarged and haemorrhagic, especially the pre-scapular lymph nodes. The most striking microscopic lesion was a necrotizing vasculitis that often started from the deep cutaneous plexus with abundant surrounding infiltrates of predominate large histiocytes and fibroblasts (Figure 2A). The histiocytes frequently contained a large, prominent eosinophilic or amphophilic, intracytoplasmic inclusion bodies and had marginated chromatin (Figure 2B). Other lesions included epidermal ballooning degeneration, subcorneal vesicles, and intracytoplasmic inclusion bodies within keratinocytes.

**Figure 2.**
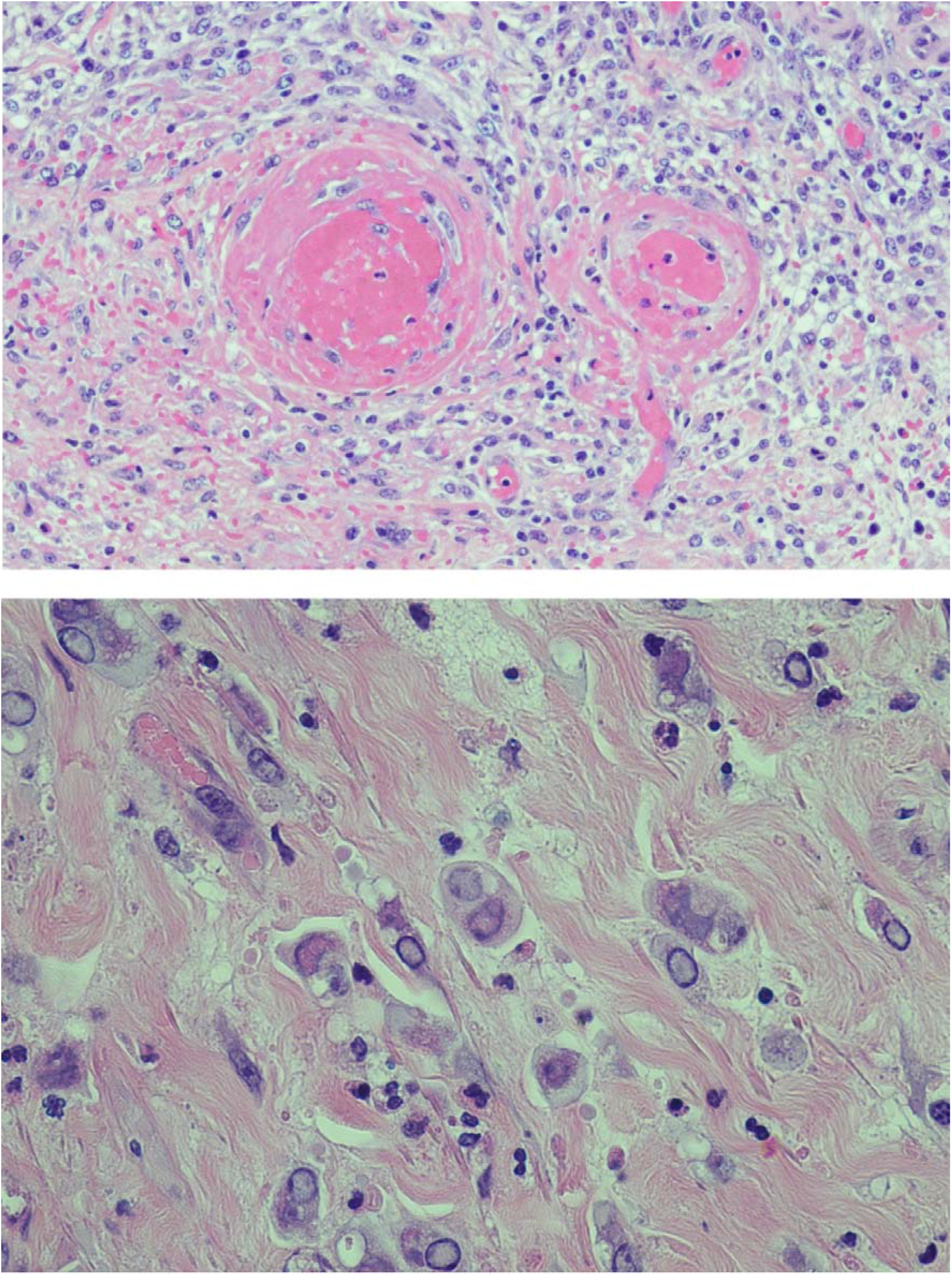
LSD causes marked, multifocal, necrotising and histiocytic dermatitis with (A) necrotising vasculitis and (B) intracytoplasmic inclusion bodies.

The clinical features of LSD caused by LSDV/HongKong/2020 strain in the field appear similar to those reported from outbreaks in south east Europe, Africa and Asia. Previous studies have reported morbidities from 1-20% and mortality of 1% or less [4–7, 32] which are consistent with the morbidity and mortality estimates of 20-30% and 0% reported in the outbreak in Hong Kong. Challenge studies carried out under experimental conditions will be required to compare in more detail the virulence of LSDV/HongKong/2020 with wildtype and KSGP strains.

Tissue samples, including subcutaneous nodules and enlarged lymph nodes were submitted to the OIE Reference Laboratory for LSD at The Pirbright Institute, UK. The capripoxvirus gene LSDV074 was detected in EDTA blood, tissue and swab samples (n=12), confirming the clinical diagnosis. LSDV was then speciated in each sample using the differentiation assay targeting LSDV011. Finally, LSDV Neethling vaccine strain was identified using the vaccine-specific assay targeting LSDV008. Both serum samples were positive for capripoxvirus antibodies. The RP030 full gene sequences were amplified and compared to capripoxvirus reference strains and identified as LSDV with the closest relatives being Neethling vaccine strains (AF409138, KX764643, KX764644, KX764645).

LSDV was isolated from skin samples taken from all three animals and named LSDV/HongKong/2020/01 to 03. Viral DNA from LSDV/HongKong/2020/01 was purified, sequenced and the genome assembled. The genome sequence has been named LSDV/HongKong/2020 and deposited in Genbank [MW732649.1]. The genome of LSDV/HongKong/2020 was compared to published genomes of other LSDV strains (Table 1). Comparison of the 90 core genes that are conserved in all chordopoxviruses [31] revealed that LSDV/HongKong/2020 was most closely related to Neethling strains of LSDV and the Saratov/2017 strain, and more distantly related to LSDV isolates from the Middle East, Europe and neighbouring regions (Figure 3). This reveals that the LSD outbreak in Hong Kong in 2020 and the LSD outbreaks in south east Europe in 2015-2017 were caused by different strains of LSDV.

**Figure 3.**
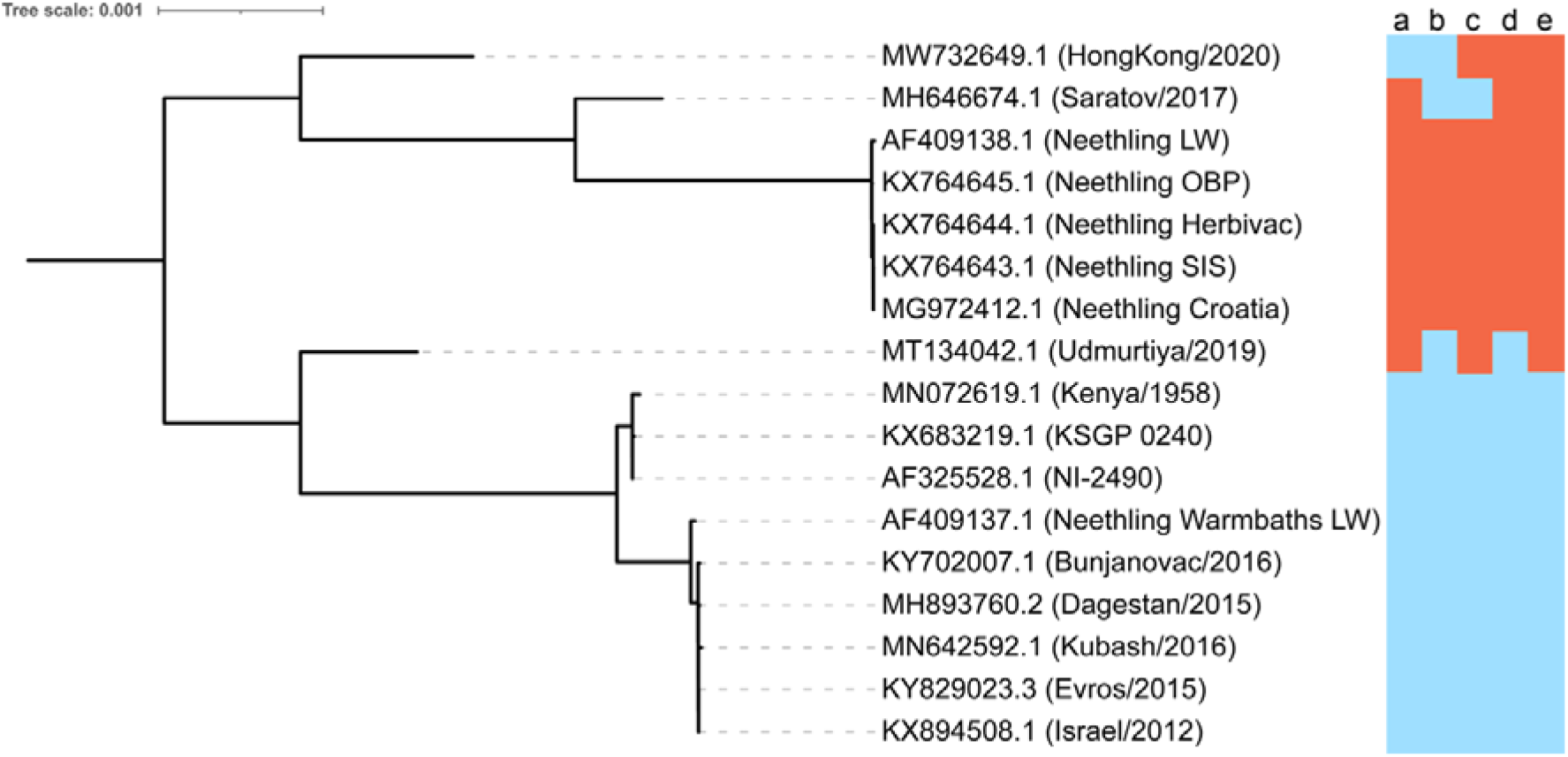
Core gene phylogenetic tree. The phylogenetic tree was built using the core genes (including intergenic regions between consecutive core genes) using RAxML and midpoint rooted. For accessory genes, isolates that grouped with either vaccine strains or wildtype strains in the network analysis are hightlighted with red and blue respectively (panel on right). Different patterns of grouping are seen for 39 accessory genes: **a)** LD006, LD007, LD143, LD148, **b)** LD030, LD133, LD144, LD145, LD147, **c)** LD009, LD135, LD136, LD137, LD138, **d)** LD015, LD017, LD034, LD035, LD127, LD141, LD142, LD146, LD150, LD151, **e)** LD008, LD010, LD018, LD020, LD021, LD023, LD066, LD067, LD122, LD125, LD126, LD128, LD129, LD130, LD152.

More detailed analysis of the accessory genes of the LSDV genomes revealed 39 genes that formed separate groupings in the network analysis when a more stringent kNN filter was applied. In Figure 3 those that grouped with the vaccine strain Neethling LW are labelled red, and those that grouped with the wildtype LSDV strain 155920/2012 are labelled blue. Five different patterns of grouping across the LSDV genomes were noted (a-e). Regardless of the genes, most wildtype and KSGP isolates grouped with wildtype LSDV strain 155920/2012, whereas the Neethling strains grouped with Neethling LW. However three LSDV strains each showed unique patterns of these accessory gene alleles – Saratov/2017, Udmurtiya/2019 and LSDV/HongKong/2020. These three strains shared higher similarity to the wildtype LSDV strain 155920/2012 for some accessory genes and higher similarity to the Neethling LW vaccine strain for other accessory genes.

The implications of the identification of the novel LSDV/HongKong/2020 strain on the diagnosis and control of LSD were considered. Diagnostic qPCR assays to differentiate between infected and vaccinated animals should be used with caution, as in this instance we were unable to use the LSDV vaccine-like assay [15] to differentiate between LSDV/HongKong/2020 and the Neethling vaccine strain due to sequence similarity within the amplified region. Redesigning these assays will be required to enable differentiation of this newly circulating strain from Neethling strain vaccines. Currently available live-attenuated LSDV vaccines are likely to provide protection against LSDV/HongKong/2020 since poxviruses are known to provide broad within-genus protection, however vaccination and challenge studies should be undertaken to confirm this.

This study has shown the value of sequencing the whole genome of LSDV rather than a small number of individual genes. We recommend carrying out WGS of new isolates of LSDV whenever possible and making the data available promptly to allow rapid identification of new LSDV strains and facilitate understanding of LSDV epidemics.

## Acknowledgements

The Pirbright Institute is funded by the Department for Environment, Food and Rural Affairs [grant code: SE26081] and the Biotechnology and Biological Sciences Research Council (BBSRC) [grant code: BB/R008833/1]. IRH and PMB were supported by BBSRC strategic funding to the Pirbright Institute [grant codes: BBS/E/I/00007036, BBS/E/I/00007037 and BBS/E/I/00007039]. BS, AL, SL and PMB are supported by a BBSRC Institute Strategic Programme grant to Roslin Institute [grant code: BBS/E/D/20002173], and SL is additionally supported by Scottish Government Rural and Environment Science and Analytical Services Division as part of Centre of Expertise on Animal Disease Outbreaks (EPIC).

## Ethical guidelines

The authors confirm that the ethical policies of the journal, as noted on the journal’s author guidelines page, have been adhered to. No ethical approval was required for this study.

## Author contributions

J.F., B.S., I.R.H., A.L., S.L., C.B, and P.M.B. conceptualised the study; J.C. carried out the clinical analyses; B.S., I.R.H., M.A., A.C., G.F., S.K., A.T., C.J.B., J.C., P.P., N.P. carried out the laboratory assays; A.D.F. carried out the necropsy and histopathology; J.F, S.K., B.S., I.R.H., A.L, S.L, and P.M.B. analysed the data; P.M.B. drafted the paper. All authors discussed the results, commented on the manuscript, and gave final approval of the version to be published.

